# Choosing the right label for single-molecule tracking in live bacteria: Side-by-side comparison of photoactivatable fluorescent protein and Halo tag dyes

**DOI:** 10.1101/381665

**Authors:** Nehir Banaz, Jarno Mäkelä, Stephan Uphoff

**Author notes:** These authors contributed equally to the work (Nehir Banaz, Jarno Mäkelä).

## Abstract

Visualizing and quantifying molecular motion and interactions inside living cells provides crucial insight into the mechanisms underlying cell function. This has been achieved by super-resolution localization microscopy and single-molecule tracking in conjunction with photoactivatable fluorescent proteins. An alternative labelling approach relies on genetically-encoded protein tags with cell-permeable fluorescent ligands which are brighter and less prone to photobleaching than fluorescent proteins but require a laborious labelling process. Either labelling method is associated with significant advantages and disadvantages that should be taken into consideration depending on the microscopy experiment planned. Here, we describe an optimised procedure for labelling Halo-tagged proteins in live *Escherichia coli* cells. We provide a side-by-side comparison of Halo tag with different fluorescent ligands against the popular photoactivatable fluorescent protein PAmCherry. Using test proteins with different intracellular dynamics, we evaluated fluorescence intensity, background, photostability, and single-molecule localization and tracking results. Capitalising on the brightness and extended spectral range of fluorescent Halo ligands, we also demonstrate high-speed and dual-colour single-molecule tracking.

## Introduction

The combination of super-resolution localization microscopy with single-particle tracking has proven very powerful as it allows direct visualization of the activity of thousands of proteins inside individual living cells. The localizations, mobility, and movement patterns of each molecule report on the molecular interactions and reactions in real-time. These data provide important quantitative information about those interactions, such as the diffusion coefficients of the reactants, dissociation constants, and the spatial distribution of interaction sites relative to other landmarks in the cell. Various implementations of this approach have provided crucial insights into many fundamental molecular processes in prokaryotic and eukaryotic cells [1–7].

Super-resolution localization microscopy is based on the sequential detection and localization of individual fluorescent molecules over the course of a movie [8]. This demands fluorescent labels with certain properties [9–11], most notably (i) means for specific labelling of target proteins ideally in a genetically-encoded manner, (ii) sufficient brightness for detection and accurate localization of single molecules, and (iii) the ability to switch fluorophores between fluorescent and non-fluorescent states so that only a fraction of molecules is visible at any time. For single-molecule tracking, photostability is also an important requirement as it limits the observation time per molecule.

Live-cell localization microscopy was enabled by the development of photoactivatable fluorescent proteins (PA-FPs) such as PA-GFP, PAmCherry, mEOS, and other variants [10–12]. In fixed and permeabilized cells, antibody staining permits the use of synthetic dyes which are generally brighter and more photostable than PA-FPs. Fixed samples can be imaged in special buffers that induce reversible photoswitching of many types of synthetic fluorophores [13,14]. More recently, genetically encoded protein tags have been developed for labelling with synthetic dyes that permeate live cells [15]. Especially Halo tag [16], SNAP tag [17], and CLIP tag [18] are becoming increasingly popular in fluorescence microscopy. In principle, these tags combine the advantages of genetically encoded FPs with the superior brightness and photostability of synthetic dyes. Because of this, they have found many successful applications in both eukaryotes and prokaryotes, using super-resolution microscopy [19,20], single-molecule imaging [21,22], and single-molecule tracking techniques [5,7,23]. Halo tag is a monomeric 33 kDa protein that was engineered from a bacterial hydrolase enzyme to form a covalent bond with a chloroalkane linker of a ligand. This reaction is rapid and irreversible under physiological intracellular conditions [16]. Promega have commercialised the system and supply a range of fluorescent ligands. A new palette of bright cell-permeable fluorescent Halo and SNAP tag ligands adds to the appeal of this labelling approach [24]. However, there are also significant disadvantages and considerations concerning these tags, as the extra labelling step introduces additional experimental complexity and labour, and labelling specificity is not guaranteed. Nevertheless, the separation of protein expression and fluorescence labelling provides another means of experimental control and opportunities for new microscopy-based assays.

Many factors need to be considered when choosing the optimal labelling strategy for any microscopy experiment, and the additional temporal dimension in single-molecule tracking presents further challenges. To facilitate the decision, we evaluated the Halo tag with different fluorescent ligands (TMR, JF549, PA-JF549, JF646) in a detailed side-by-side comparison against PAmCherry, a popular PA-FP [12]. We optimised the protocol for Halo tag labelling, tested labelling specificity and background, quantified the brightness and photostability of the labels, and performed tracking experiments on several different fusion proteins. We compile these assessments to provide a practical guide for choosing the most suitable label for single-molecule tracking applications in live *E. coli* cells.

### Fluorescent labelling of intracellular Halo tag proteins

To evaluate the performance of Halo tag and PAmCherry we chose test proteins that had previously been characterised. MukB is a Structural Maintenance of Chromosomes (SMC) protein that acts in chromosome segregation together with accessory proteins MukE and MukF at a copy number of ~200 molecules per cell [25]. Tracking MukB-PAmCherry revealed the presence of mobile and immobile SMC complexes that accumulated in distinct foci inside cells [26]. These foci appear to position the chromosome replication origin and subsequently aid in the segregation of the replicated origins into the two daughter cells [27]. As a second test protein with different intracellular mobility and localization, we chose DNA polymerase I (PolI), a monomeric protein that is much smaller than the SMC complex and present at ~400 copies per cell [28]. PolI performs DNA synthesis during the processing of Okazaki fragments and in DNA excision repair pathways [29]. Following DNA damage treatment, PolI-PAmCherry molecules become transiently immobilised at DNA repair sites [28]. To complement the existing PAmCherry fusions of MukB and PolI [26,28], we generated endogenous C-terminal translational Halo tag fusions using the Lambda Red recombination technique [30]. We detected no signs of any growth defects that would indicate impairment of the function of MukB or PolI by the Halo fusions. Imaging was performed on a custom-built Total Internal Reflection Fluorescence (TIRF) microscope equipped with an electron-multiplying CCD camera [31]. To image cytoplasmic proteins within ~1 µm thick *E. coli* cells, laser excitation was adjusted to highly inclined illumination mode [32].

We optimized a procedure for labelling Halo tag in live *E. coli* with the commercially available dye TMR (Fig. 1). Briefly, we grew cell cultures expressing PolI-Halo to early exponential phase at 37°C in M9 minimal growth medium containing glucose and amino acids. 1 ml of culture was concentrated by centrifugation, resuspended in 100 µl of growth medium and incubated with 2.5 µM TMR dye at 25°C for 30 min. Removal of unbound dye was crucial to minimize fluorescent background for robust detection of single molecules inside cells. This is especially important for imaging bacteria directly on the surface of the coverslip where fluorescent particles adhere. We optimized the washing procedure by measuring the background over multiple centrifugation and resuspension cycles (Fig. 2a). After each wash, a small amount of cell suspension was immobilized on an agarose gel pad sandwiched between two microscope coverslips and imaged using 561 nm laser excitation. Intracellular fluorescence became visible above the background noise after 2 – 3 washes, but substantial background remained both on the coverslip surface and as diffuse background within the agarose pad. Therefore, to facilitate release of unbound dye and to recover the cells from the prolonged centrifugation, we performed a 4^th^ wash and incubated the diluted labelled culture at 37°C for 30 min in a shaking incubator, followed by a final 5^th^ wash. This removed the fluorescent background almost entirely. The entire labelling, washing, and recovery procedure takes approximately 90 min. Note that cell growth and division during the recovery dilutes the labelled Halo proteins exponentially while new unlabelled Halo is synthesised.

**Figure 1.**
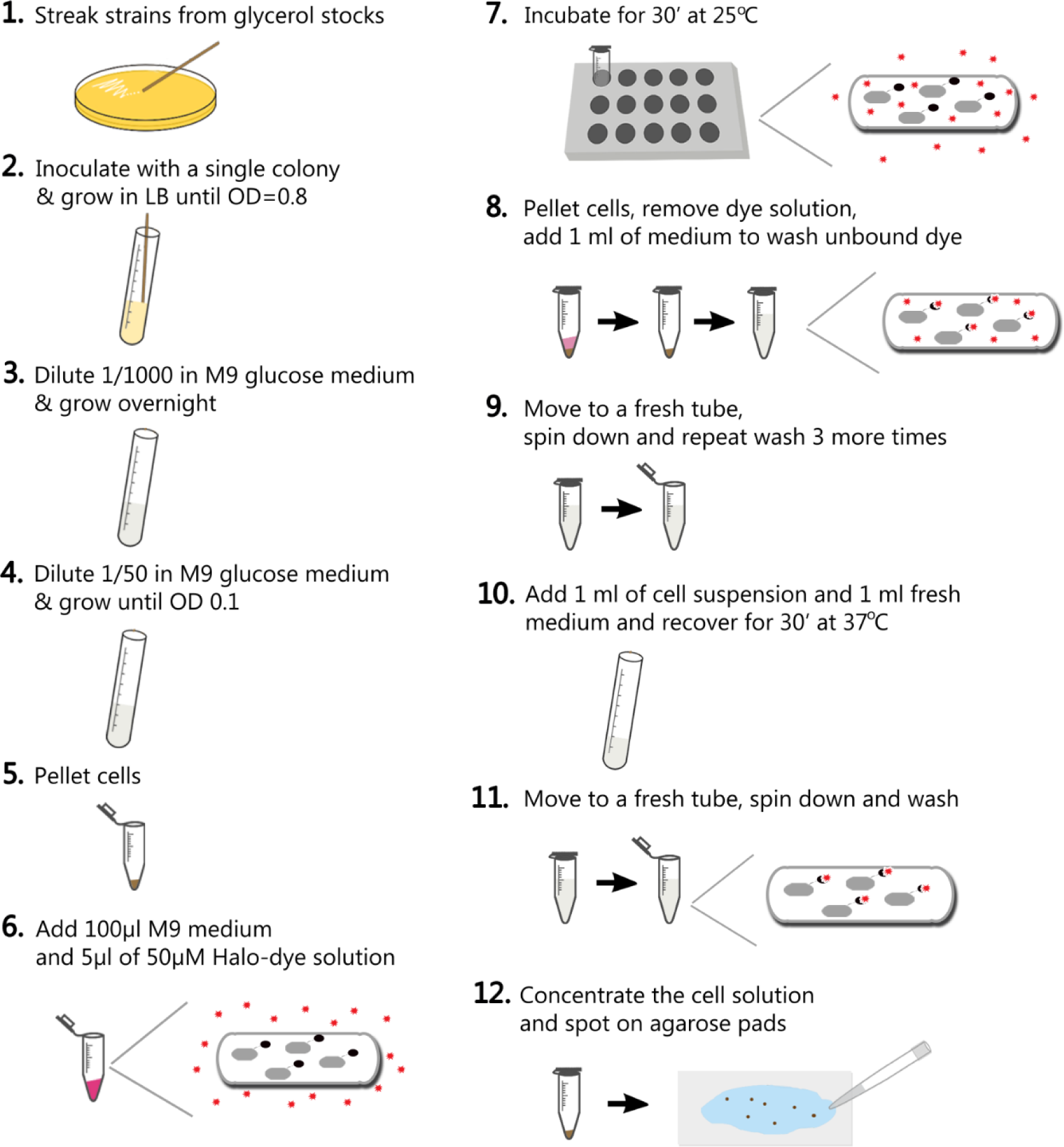
Procedure for labelling Halo tag with a fluorescent ligand in live *E. coli* cells.

**Figure 2.**
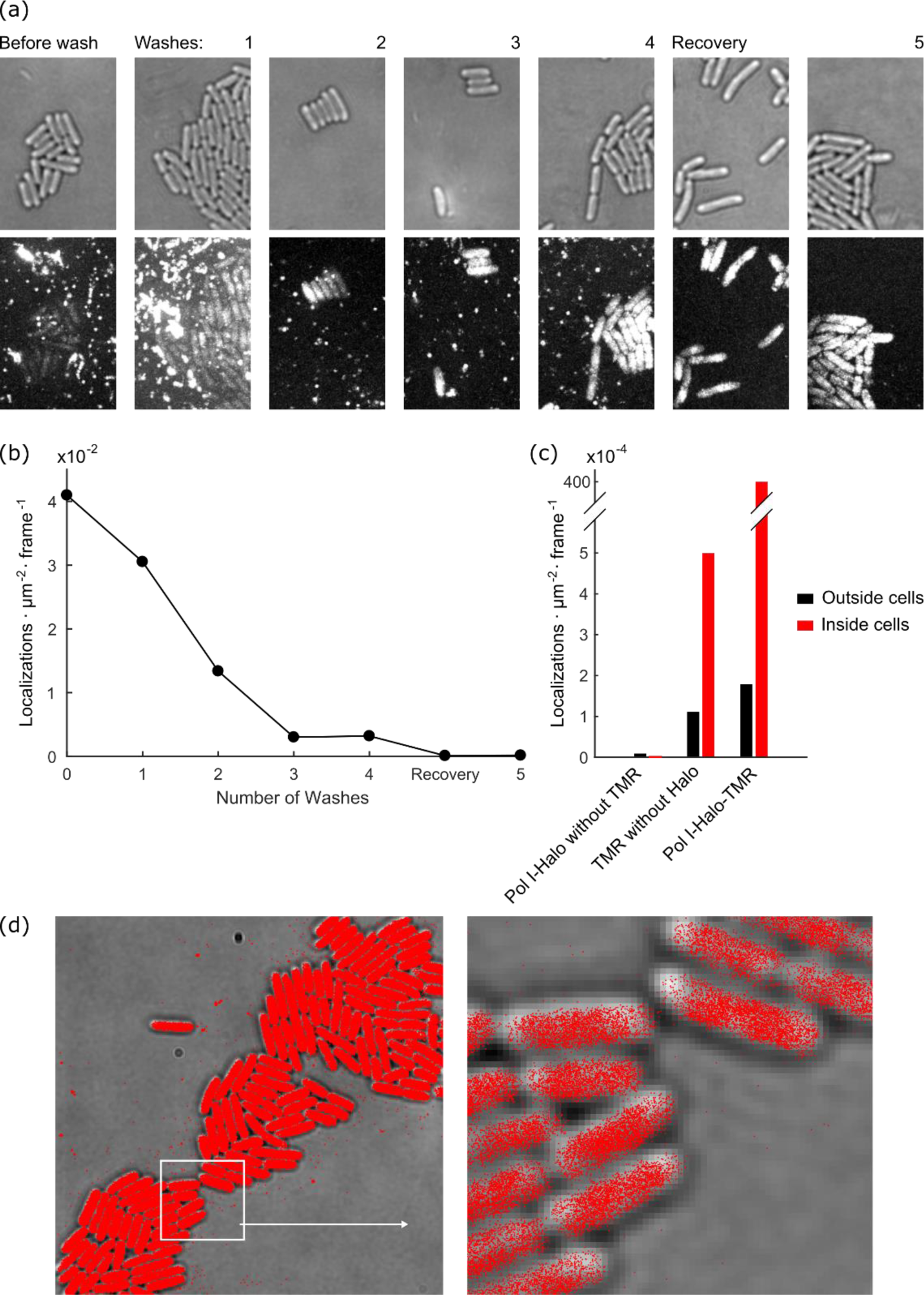
Removal of fluorescent background after Halo tag labelling. (a) *E. coli* expressing PolI-Halo-TMR were labelled and a super-resolution movie of 15,000 frames was recorded after each washing step to remove unbound dye. Transmitted light images show cell outlines (top). Fluorescence images show maximum intensity projections from the super-resolution movie (bottom, identical grey scaling across images for all washes). (b) Frequency of background localizations outside cells during the washing procedure. The frequency represents the average number of localizations detected per area per frame. (c) Frequency of localizations outside cells and inside cells: (i) for cells without TMR labelling, (ii) for cells that do not express Halo tag but were labelled with TMR (5 washes), (iii) for cells expressing PolI-Halo that were labelled with TMR (5 washes). (d) Map of PolI-Halo-TMR localizations (5 washes) plotted on transmitted light snapshot. The boxed area is shown magnified.

To quantify how the background affects single-molecule localization microscopy, we acquired movies and performed localization analysis as described below and in our previous work [33]. The frequency of background localizations outside cells decreased with each washing cycle from an average of 4.1∙10^-^ 2 localizations∙µm^-2^∙frame^-1^ without washing to 1.7∙10^-4^ localizations∙µm^-2^∙frame^-1^ after 5 washes (Fig. 2b). The remaining background after the washing procedure was much lower than the desired signal from PolI-Halo-TMR inside cells (average frequency of 4∙10^-2^ localizations∙µm^-2^∙frame^-1^, Fig. 2c-d). To assess the level of non-specific labelling, we measured the localization frequency of TMR inside cells that do not express Halo tag. The localization frequency was approximately 5-fold above the background outside cells, but more than 2 orders of magnitude lower than the specific localizations of PolI-Halo-TMR (Fig. 2c). Furthermore, the localization frequency inside cells without any TMR labelling was as low as 2∙10^-6^ localizations∙µm^-2^∙frame^-1^ (Fig. 2c). Therefore, the optimised labelling and washing procedure, produces reliable single-molecule localization data for Halo-TMR in live *E. coli* cells with little non-specific background (Fig. 2d).

### Single-molecule localization microscopy of Halo tag and PAmCherry fusions

To test the utility of the Halo tag for imaging intracellular protein localization, we recorded snapshots of MukB-Halo-TMR using a standard Nikon epifluorescence microscope. Comparison with a fusion of MukB to the conventional fluorescent protein mCherry shows that both labelling strategies reproduce the characteristic foci of MukB [26] with similar image quality (Fig. 3a).

**Figure 3.**
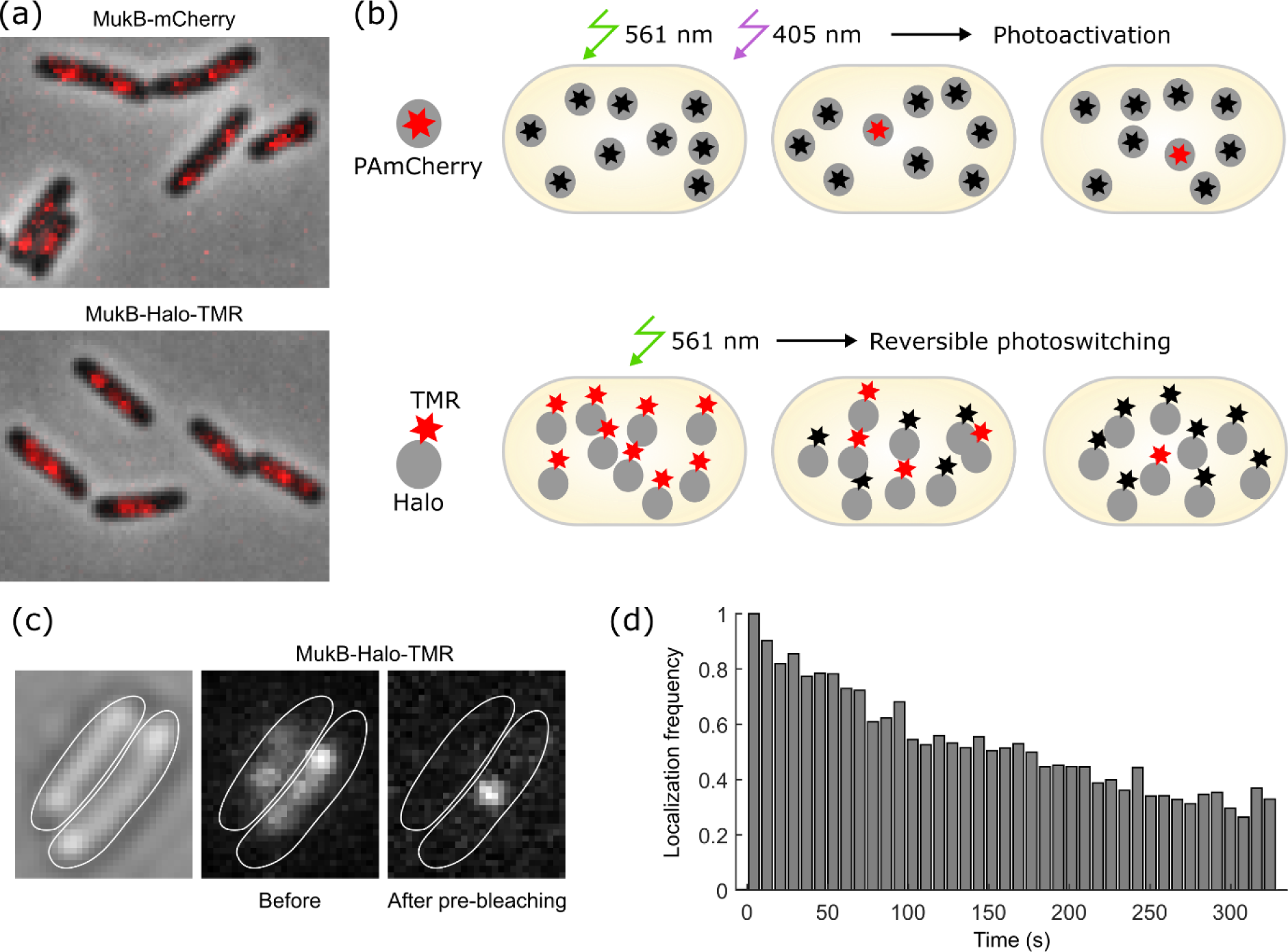
Microscopy data acquisition using Halo tag. (a) Snapshots of MukB-Halo-TMR and MukB-mCherry recorded with identical settings on an epifluorescence microscope (red: fluorescence, grey: phase contrast). (b) Principle of data acquisition for single-molecule localization microscopy using PAmCherry or Halo-TMR labels. (c) Snapshots showing MukB-Halo-TMR intensity before and after pre-bleaching. (d) Frequency of MukB-Halo-TMR localizations over the course of a movie at constant 561 nm excitation (0.2 kW/cm^2^), normalised to the initial frequency.

The procedure for recording single-molecule localization or tracking data with Halo-TMR differs from PAmCherry. PAmCherry is initially in a non-fluorescent state and becomes converted by 405 nm illumination to the fluorescent state which can be excited at 561 nm [10,12]. Consequently, the sample appears dark at the beginning of the experiment apart from a small number of spontaneously activated molecules (Fig. 3b). On the one hand, this can be a technical inconvenience, as the absence of any signal makes it harder to find regions of interest and to adjust the focus. On the other hand, illumination at 561 nm without 405 nm photoactivation allows bleaching any background fluorescence before data acquisition. Simultaneous 405 nm and 561 nm illumination during the acquisition causes continuous activation and bleaching of PAmCherry molecules for sequential localization of individual molecules (Fig. 3b). Adjusting the 405 nm intensity provides accurate control over the density of fluorescent dyes at any time to facilitate single-molecule detection. Because most PAmCherry molecules bleach irreversibly after one photoactivation cycle, the decay in the density of fluorescent molecules can be counteracted by gradually increasing the 405 nm intensity during an acquisition.

Contrary to PAmCherry, TMR-labelled Halo fusions are initially in the fluorescent state (Fig. 3b,c). Therefore, pre-bleaching is required before single-molecule imaging, but this can be useful for capturing a conventional epifluorescence image as a low-resolution reference (Fig. 3d). Under continuous illumination at 561 nm, TMR showed both photobleaching and reversible photoswitching within live cells. Initially, the number of visible fluorophores decayed rapidly due to reversible deactivation of the dye (Fig. 3c), followed by a long period of spontaneous photoswitching (Fig. 3d). The frequency of localisations decreased very slowly over the course of a movie because of photobleaching, with ~40% of fluorophores remaining even after 5 minutes of continuous imaging (Fig. 3d). Importantly, imaging was performed in M9 minimal medium containing only supplements for cell growth (glucose, amino acids, and thiamine). There was no requirement for special buffer additives to induce photoswitching (e.g. oxygen scavenging and reducing agents) because the reducing environment inside cells promotes the photoinduced formation of a dark radical anion state that is recovered by molecular oxygen [34]. The lifetime of the reversible dark state of TMR was sufficiently long such that only a sparse subset of molecules appeared fluorescent at any time, which is an important requirement for accurate single-molecule localization and tracking. Compared to PAmCherry, the density of fluorescent TMR molecules cannot be precisely controlled at this stage in the experiment. However, if the density is too high, the initial photobleaching can be prolonged before data acquisition or substoichiometric labelling can be achieved by using a lower concentration of the dye.

We generally detected more localizations from Halo-TMR compared to PAmCherry under the same acquisition settings. This was mostly due to the repeated photoswitching cycles of TMR as opposed to irreversible activation and bleaching of PAmCherry. Although repeated photoswitching of TMR provides more localization data from the same number of molecules per cell, this may complicate quantitative analysis of the localization data. In particular, blinking of single molecules produces apparent clusters of localizations, which are difficult to distinguish from genuine molecular assemblies [35]. This issue also affects PA-FPs, albeit to a lesser extent than photoswitchable dyes [36]. Several data analysis strategies have been reported to correct for blinking effects in molecule counting and clustering analysis [35–39].

The average intensity of single MukB-Halo-TMR molecules was approximately 2-fold higher than MukB-PAmCherry under the same imaging conditions (Fig. 4a). Notably, fluorescence brightness saturated with increasing 561 nm excitation intensity for both dyes, while the background shot noise continues to increase (Fig. 4b). This creates a peak in the signal to noise ratio at ~0.2 kW/cm^2^ 561 nm intensity (Fig. 4c). We also tested the Halo ligand JF549, which is a derivative of TMR with improved brightness and photostability [24]. We achieved excellent labelling of MukB-Halo-JF549 using the same protocol as for TMR, and found that it showed similar photoswitching behaviour. JF549 was marginally brighter than TMR at the optimal excitation intensity of 0.2 kW/cm^2^ (Fig. 4a-c). Photoactivatable Halo tag ligands have also been developed [40]. We tested one such dye, PA-JF549, but found that it did not label MukB-Halo efficiently using our protocol. Only ~1% of cells showed intracellular fluorescence, likely because the dye does not pass the *E. coli* cell wall.

**Figure 4.**
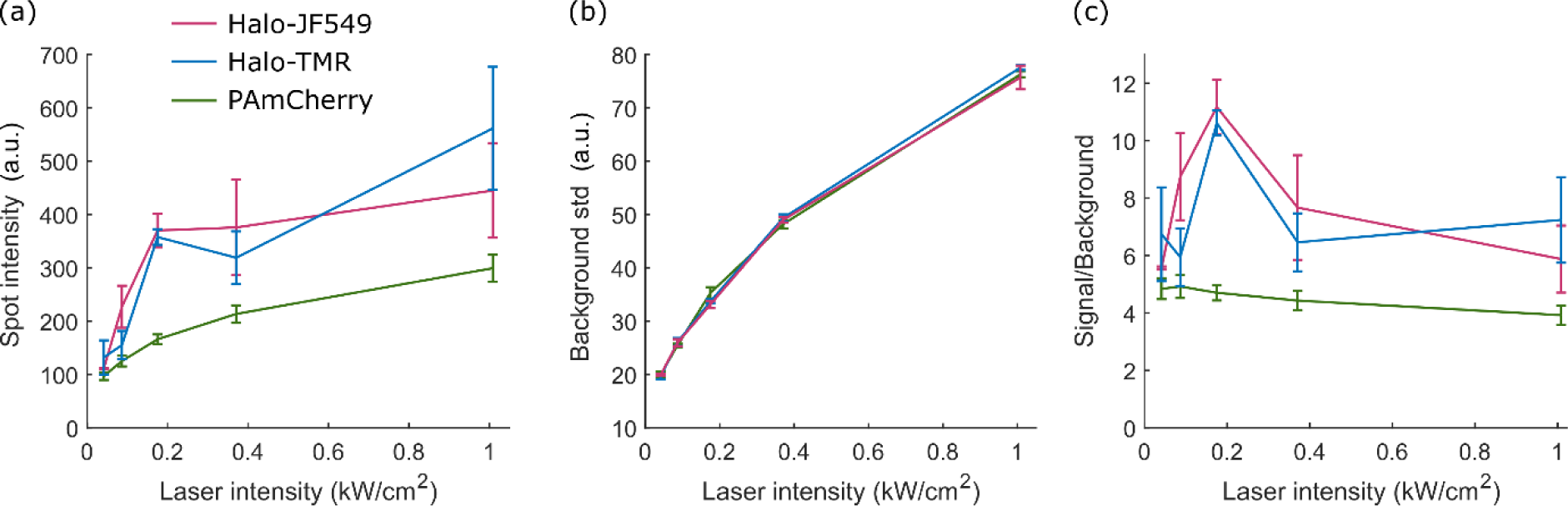
Fluorescence intensity comparison of PAmCherry, Halo-TMR, and Halo-JF549. (a) Fluorescence spot intensity (a.u.) of single MukB fusion proteins tagged with the indicated fluorophores as a function of 561 nm excitation intensity. (b) Standard deviation of the background noise (a.u.) as a function of 561 nm excitation intensity. (c) Signal to background ratio for the data in a-b.

### Single-molecule tracking of Halo tag and PAmCherry fusions

The use of photoactivatable or photoswitchable fluorophores generalizes single-particle tracking, such that a very large number of trajectories can be recorded per cell, irrespective of the protein expression level or labelling density [41]. A major appeal of single-molecule tracking experiments is the ability to directly observe and quantify molecular interactions. PA-FPs have been used successfully for this purpose [2,4,6,42]. However, low brightness and photostability of these fluorophores have been major limitations. Fluorophore brightness not only determines the spatial resolution, but also sets a limit on the temporal resolution at which molecules can be tracked. Measuring protein motion in the bacterial cytoplasm requires imaging at high frame rates of typically 10-1000 frames/s. Because photobleaching restricts the observation time per molecule, it has been challenging to record long-lived, rare, or transient molecular events. We thus tested whether the enhanced photophysical characteristics of fluorescent Halo tag ligands enable faster and longer observations, which would open avenues for new biological applications.

Using a tracking algorithm, 2D localizations were linked to tracks if they appeared within a radius of 768 nm (8 pixels) in consecutive frames [33]. To reconnect tracks with single missed localizations, we used a memory parameter of 1 frame. An apparent diffusion coefficient D* = MSD/4Δt was calculated from the mean squared displacement (MSD) between consecutive localizations of individual tracks with Δt = 15.48 ms. For our purposes, D* serves as a relative measure of mobility but does not represent the accurate diffusion coefficient of a molecule because of several systematic biases, such as motion confinement, motion blurring, and localization error. If required, these biases can be estimated and corrected for according to previously described procedures [28,33]. The distribution of D* values for MukB-Halo-TMR showed a mixture of immobile (D* ~ 0.05 µm^2^/s) and slowly diffusing (D* ~ 0.3 µm^2^/s) molecules (Fig. 5a). By classifying tracks according to a threshold of D* = 0.15 µm^2^/s, it is evident that stationary molecules localized in clusters, whereas diffusing molecules moved randomly within the nucleoid (Fig. 5b). These results match our observations for MukB-PAmCherry (Fig. 5a-b), and reproduce previous findings [26].

**Figure 5.**
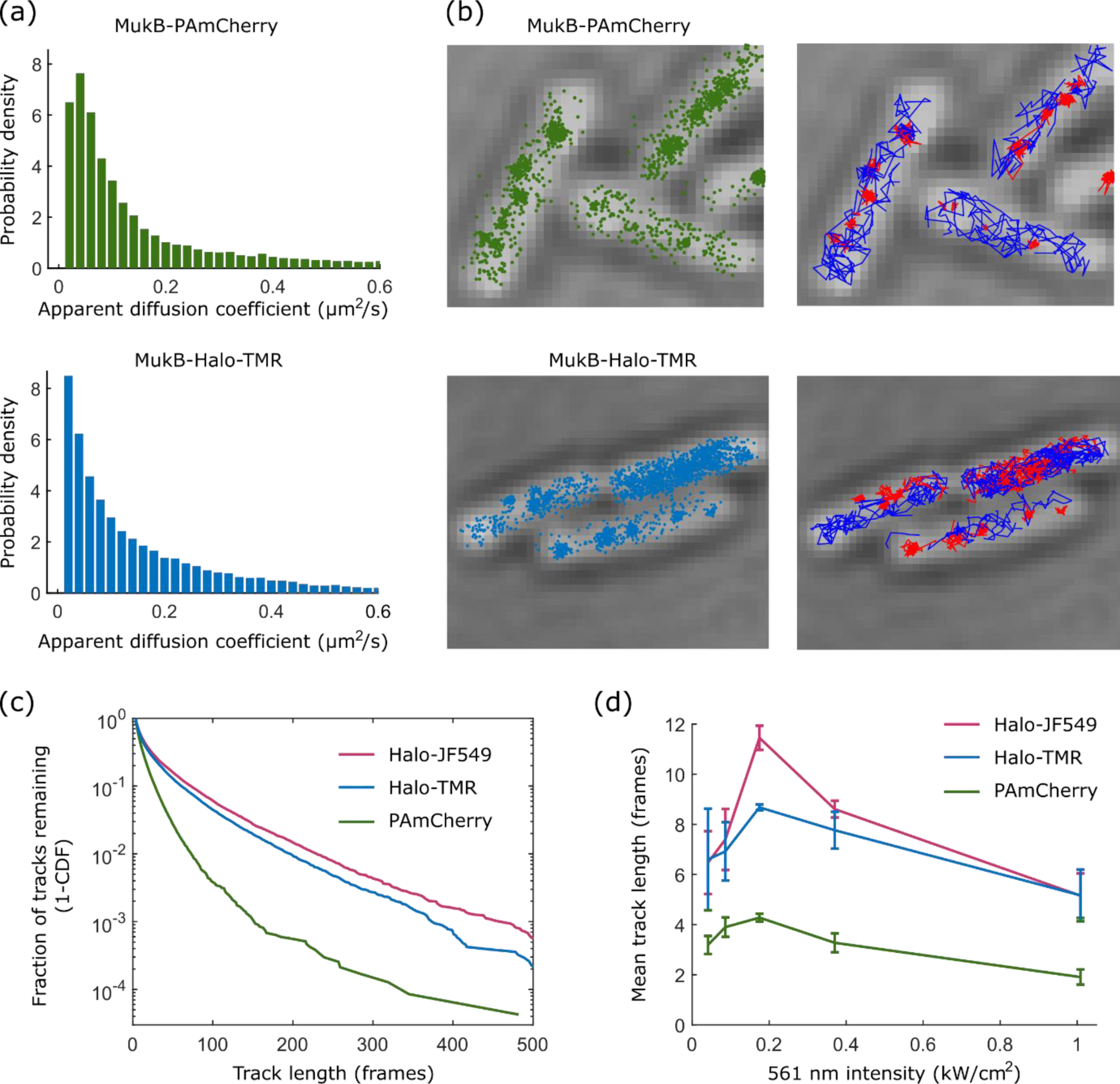
Application of PAmCherry and Halo tag labels for single-molecule tracking of MukB. (a) Distributions of the apparent diffusion coefficient for MukB-PAmCherry and MukB-Halo-TMR. (b) Localisations and tracks of MukB-PAmCherry and MukB-Halo-TMR. Stationary tracks with D* < 0.15 µm2/s are shown in red, mobile tracks in blue. (c) Track length distributions for MukB labelled with PAmCherry, Halo-TMR, and Halo-JF549. (d) Mean track length as a function of 561 nm excitation intensity.

Next, we analysed the duration of single-molecule tracks for MukB-PAmCherry, MukB-Halo-TMR, and MukB-Halo-JF549. For PAmCherry, trajectories are terminated mainly by photobleaching, whereas both reversible photoswitching and photobleaching occur for TMR and JF549. Track durations were significantly longer for the Halo dyes compared to PAmCherry (Fig. 5c). At optimal excitation intensity (0.2 kW/cm^2^ 561 nm, Fig. 5d), the average track durations were 4.3, 8.7, and 11.5 frames for PAmCherry, TMR, and JF549, respectively. The distribution of track durations shows a long tail with 4.5% and 6.1% of TMR and JF549 molecules lasting for more than 100 frames, compared to 0.4% for PAmCherry. This enables direct observations of protein function within living cells over much longer time scales than previously possible.

We used the PolI fusions to test if the Halo tag is also suitable for the measurement of transient molecular interactions. PolI is an essential factor for the repair of DNA base damage, filling DNA gaps that are generated by the base excision repair machinery [29]. The majority of PolI-Halo-TMR molecules were diffusing with D* ~ 0.7 µm^2^/s in cells before DNA damage treatment (Fig. 6a). To monitor the DNA damage response, we first labelled PolI-Halo with TMR, and subsequently treated cells on agarose pads containing the DNA damaging agent methyl methanesulfonate (MMS). We observed transient immobilization of individual PolI molecules after MMS exposure for PAmCherry and Halo-TMR fusions (Fig. 6b), as previously shown [28]. A global shift in the mobility of PolI-Halo-TMR molecules occurred after 45 min of MMS treatment (Fig. 6c), which we attribute to the compaction of the nucleoid in response to DNA damage [28]. This effect was stronger for Halo compared to PAmCherry, which likely reflects the differences in the labelling procedure. The high density of cells during TMR labelling and washing reflects the conditions that cells experience in stationary growth phase, which promotes nucleoid compaction. This is an example where the Halo labelling procedure appears to affect the biological process under study.

**Figure 6.**
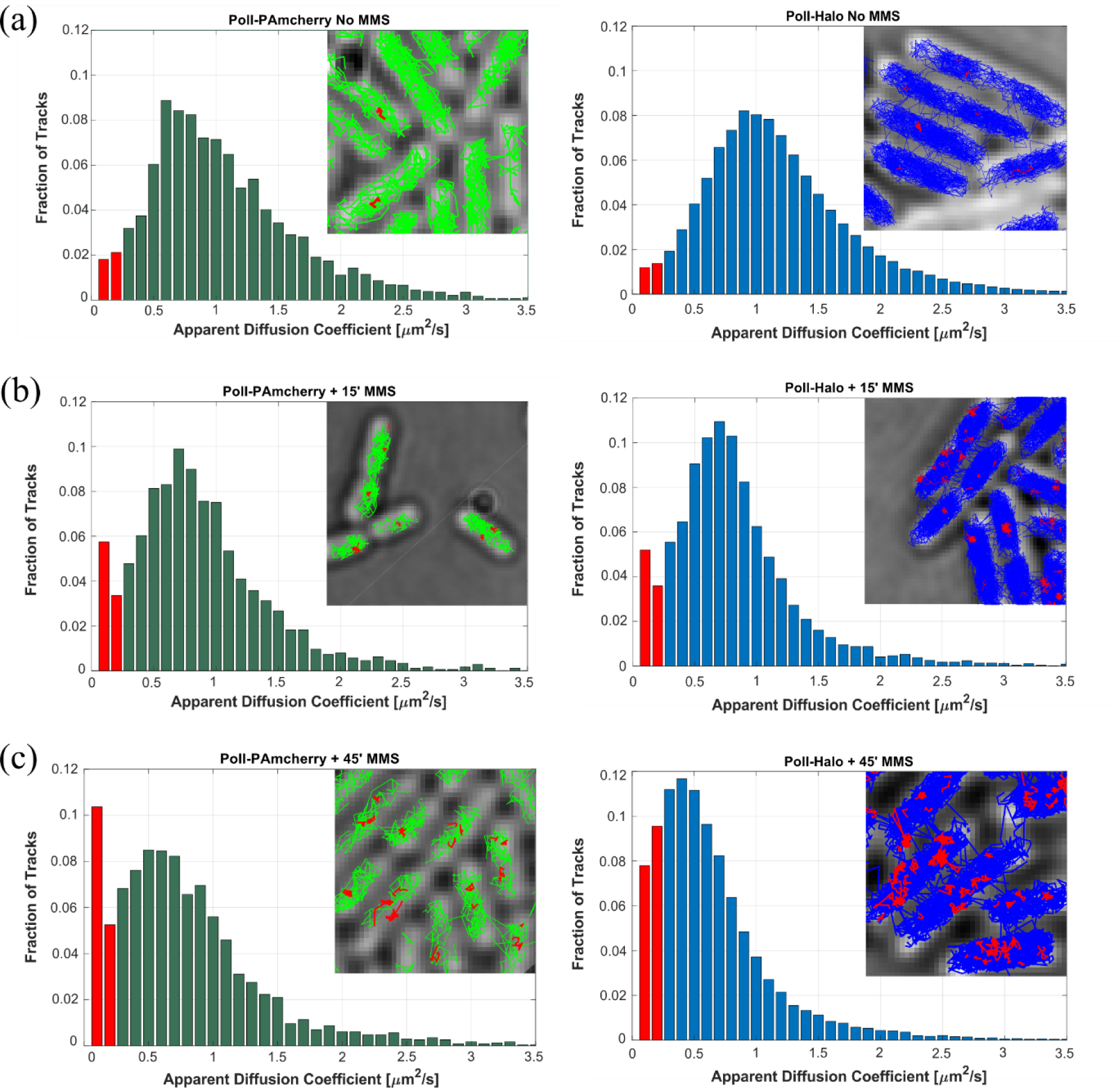
Tracking DNA repair activity of PolI. Cells with PolI-PAmCherry (left) or PolI-Halo-TMR (right) were imaged before (a) and after treatment with 100 mM methyl methanesulfonate (MMS) for 15 min (b) and 45 min (c). Immobile molecules with D* < 0.15 µm^2^/s are shown in red.

### Halo labelling enables high-speed and dual-colour tracking

To track rapidly diffusing proteins, previous studies relied on stroboscopic illumination patterns in order to reduce blurring of the fluorescent spots due to movement of molecules during the exposure [43,44]. The superior brightness of fluorescent Halo ligands may enable tracking fast molecular motion with simple continuous illumination. Avoiding stroboscopic pulses would be desirable not only for technical convenience, but also to reduce the phototoxicity assosciated with the peak excitation intensities [45]. To test the limits of high-speed single-molecule tracking, we measured unconjugated Halo molecules that were expressed from a plasmid and labelled with TMR (Fig. 7a). With continuous 561 nm illumination (0.4 kW/cm^2^) and a frame rate of 134 frames/s (7.48 ms/frame), fluorescent spots were sufficiently bright for robust localization and tracking despite residual motion blurring (Fig. 7b,c). This was in stark contrast to unconjugated PAmCherry molecules, which were almost indistinguishable from the background noise at such short exposure times. Because any protein fused to Halo should have a lower mobility than the Halo tag alone, this result demonstrates the technical feasibility of tracking even the most rapidly diffusing fusion proteins with simple continuous illumination. Stable non-specific binding of unconjugated Halo molecules was reported in eukaryotic nuclei [46]. This complicates the interpretation of the tracking data and may perturb the function of fusion proteins. Here, we observed only a small fraction of apparently immobile Halo-TMR molecules in *E. coli* (1% of all tracks, Fig. 7d), which was likely due to residual fluorescent background particles.

**Figure 7.**
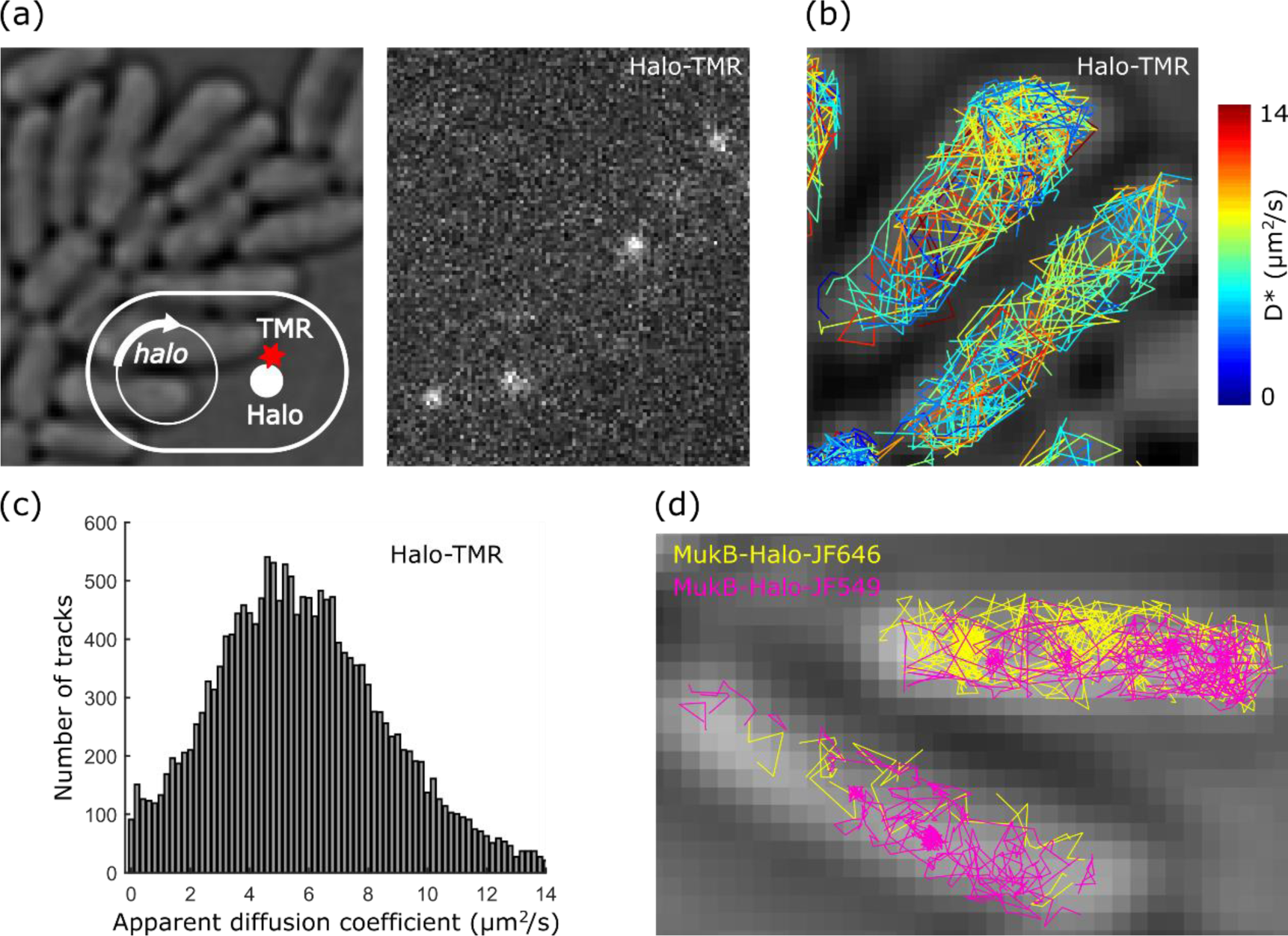
High-speed and dual-colour single-molecule tracking using Halo tag. (a) Cells expressing unconjugated Halo tag from a plasmid. Snapshot from a super-resolution movie showing Halo-TMR molecules at 7.48 ms exposure time. (b) Halo-TMR tracks in colours corresponding to the apparent diffusion coefficient. (c) Histogram of apparent diffusion coefficients for Halo-TMR. (d) Dual-colour tracking of MukB-Halo-JF646 (yellow) and MukB-Halo-JF549 (pink).

The ability to use Halo tag with a range of different ligands is a further advantage of this labelling approach. It facilitates multi-colour imaging and enables the use of fluorophores in the far-red spectrum where cellular auto-fluorescence and phototoxicity are less pronounced. We tested dual-colour tracking using Halo labelling with JF549 and JF646 dyes [24]. As a proof-of-principle, we first labelled MukB-Halo with JF646 according to the protocol in Figure 1. Following 2 hours of cell recovery, we labelled the newly synthesised MukB-Halo with JF549 using the same protocol. We imaged the two labels sequentially, recording 10,000 frames under 640 nm excitation (0.24 kW/cm^2^) followed by 10,000 frames under 561 nm excitation (0.2 kW/cm^2^). The dual-colour tracking data shows the expected diffusing and immobile MukB molecules for both labels (Fig. 7e).

### Choosing the right label for single-molecule tracking experiments

Applications that require maximum fluorophore brightness for fast tracking or high localization precision benefit from the use of protein tags with synthetic dyes. The high excitation intensities required for super-resolution microscopy and single-molecule tracking cause significant phototoxicity in cells. Especially near-UV light required for photoactivation of PA-FPs is problematic [45]. Ongoing development of synthetic fluorophores will further improve their photophysical characteristics, cell permeability, and availability. However, several important limitations need to be taken into account. The requirement for labelling and extensive washing of the fluorescent ligand may complicate applications that require precisely controlled growth conditions or cell perturbations. Prolonged time-lapse imaging benefits from the continuous synthesis and replenishment of fluorescent proteins. Improved fluorescent proteins are also reported, with extended spectral range [47], beneficial photoactivation/conversion/switching properties, reduced maturation time [48]. We close with a table showing what we consider the most important practical considerations in favour and against PA-FPs and synthetic dyes for protein tags (Table 1).

**Table 1.** Practical considerations regarding labelling approaches for single-molecule localization and tracking experiments in live bacterial cells.

|  | Advantages | Disadvantages |
| --- | --- | --- |
| **Photoactivatable fluorescent protein (e.g. PAmCherry)** | • Genetically encoded fluorophore (no additional labelling required, labelling specificity guaranteed).<br>• Independently controlled photoactivation with 405 nm excitation.<br>• Very low background noise (background can be bleached before photoactivation).<br>• Cells can be imaged during continuous growth.<br>• Protein synthesis during cell growth replenishes fluorescence.<br>• Established and well-characterised labels. | • Low brightness limits spatial and temporal resolution.<br>• Observation time per molecule is limited by rapid photobleaching.<br>• Irreversible bleaching prohibits repeated measurements of the same molecules.<br>• Limited spectral range.<br>• Phototoxicity of 405 nm illumination for photoactivation.<br>• Multi-colour tracking is limited because most PA-FPs are activated by near-UV light.<br>• Quantitative localization analysis complicated by incomplete photoactivation and long maturation times.<br>• Aggregation artifacts have been reported. |
| **Protein tag labelled with synthetic fluorophore (e.g. Halo-TMR)** | • Superior brightness provides enhanced localization precision and temporal resolution.<br>• Reduced photobleaching increases observation time per molecule.<br>• Available dyes span visible up to far-red spectrum.<br>• Compatible with conventional fluorescence imaging.<br>• Multi-colour tracking possible using orthogonal labelling of Halo, SNAP, and CLIP tags with different dyes.<br>• Reversible blinking allows repeated measurements of the same molecules.<br>• Ability to perform sequential labelling (e.g. before and after a cell perturbation). | • Additional labelling step (laborious, may interfere with other cell treatments, time-course experiments, etc)<br>• Removal of unbound dye requires extensive cell washing and centrifugation.<br>• Labelled protein is diluted with each cell division, causing loss of signal during time-lapse imaging.<br>• Pre-bleaching step required before single-molecule imaging.<br>• Limited control over photoswitching kinetics.<br>• Reversible blinking produces apparent clustering artifacts.<br>• Quantitative localization analysis complicated by reversible blinking, unknown labelling efficiency, and dilution of labelled proteins during cell growth.<br>• Not all Halo tag dyes are cell permeable.<br>• Not all fluorescent Halo ligands are commercially available and can be expensive. |

## Acknowledgments

We thank Luke Lavis for providing the Halo ligand dyes JF549, PA-JF549, and JF646, and David J Sherratt for discussions. Imaging in the Micron Advanced Microscopy Facility was funded by Wellcome Trust Strategic Award no. 107457. This work was supported by a Sir Henry Dale Fellowship jointly funded by the Wellcome Trust and the Royal Society (Grant Number 206159/Z/17/Z) and a Wellcome-Beit Prize (206159/Z/17/B) to SU.

## Author contributions

NB and JM contributed equally to the work. SU initiated the study. NB, JM, SU performed experiments and analysed the data. NB, JM, SU prepared manuscript figures. SU wrote the manuscript with input from all authors.

